## Supplementary Results 1-3 for "CRISPR-Cas is beneficial in plasmid competition, but limited by competitor toxin-antitoxin activity when horizontally transferred"

**Supplementary Results 1 – Mathematical modelling results**

Model development

Figure 1AB outlines the basic conceptual assumptions underpinning our model. Our mathematical model focusses on the two classes of co-infected cells (*CT* and *TC*), and defines transition rates from these co-infected states to singly infected states (*C* or *T*).

These transition rates are defined by plasmid segregational loss (baseline rate *s*), modified by *C-* and *T-*specific traits of CRISPR-Cas induced enhanced segregation (multiplicative factor *x*; *x* > 1 implies increased segregation rate *xs > s*) and TA-induced PSK (probability of PSK, *y*). In the event of PSK, we assume that the liberated resources (nutrients, space) benefit a kin cell weighted by parameter *f_r_*, benefit a non-kin cell (*f_i_*), or benefit no cells (e.g. due to cell dormancy versus PSK; *f_0_*), where *f_0_* + *f_i_* + *f_r_* = 1 [1]. Finally, asymmetries in parameter values between resident and invader roles are captured by subscripts *r* and *i*. Given lower plasmid copy number and lower gene expression for invading versus established plasmids [2], our default assumptions are *s_i_* > *s_r_*; *x_r_* > *x_i_* ≥ 1; 1 > *y_r_* > *y_i_* ≥ 0. Given spatial structuring so that *T* cells are enriched in neighbourhood of *TC* cells (and vice-versa for *C* near *CT* cells), we further assume 1 > *f_r_* > *f_i_* ≥ 0.

Transitions for *CT* cells (when the CRISPR-Cas plasmid is resident) are formalised as follows.

$t_{CT,C}={x_{r}s}_{i}{((1-y}_{i})+y_{i} f_{r})$	E1; for transitions to CRISPR-Cas plasmid *C.*

$t_{CT,T}=s_{r}+x_{r} s_{i} y_{i} f_{i}$	E2; for transitions to TA plasmid *T*.

Similarly, transitions for *TC* cells (when the TA plasmid is resident) are formalised below.

$t_{TC,C}=x_{i}s_{r}(\left( 1-y_{r} \right)+y_{r} f_{i})$	E3; for transitions to CRISPR-Cas plasmid *C.*

$t_{TC,T}=s_{i}+x_{i} s_{r} y_{r} f_{r}$	E4; for transitions to TA plasmid *T.*

Mathematical analysis.

Analysis of how the transition rates (equations E1-E4) change with competitive traits *x* and *y* reveals the following predictions:

(1) In the absence of PSK (*y* = 0), increasing investment in CRISPR-Cas (increasing *x*) always increases transition gains, regardless of whether the CRISPR-Cas plasmid is in a resident or invader role:

${\frac{d t_{CT,C}}{d x_{r}}|}_{y_{i}=0}=s_{i} > 0$ ; ${\frac{d t_{TC,C}}{d x_{i}}|}_{y_{r}=0}=s_{r}> 0$

(2) In the absence of CRISPR-Cas (*x* = 1), increasing investment in TA-mediated PSK (increasing *y*) always increases *T*-plasmid gains, again regardless of role:

${\frac{d t_{CT,T}}{d y_{i}}|}_{x_{r}=1}={f_{i}s}_{i} > 0$ ; ${\frac{d t_{TC,T}}{d y_{r}}|}_{x_{i}=1}={f_{r}s}_{r}> 0$

(3) When both are active (*x* > 1 and *y* > 0), an asymmetry emerges between TA and CRISPR-Cas. TA-mediated transition gains (*t_TC,T_, t_CT,T_*) are increasing functions of CRISPR-Cas activity *x* ($\frac{d t_{CT,T}}{d x_{r}}={f_{i}s}_{i} y_{i}>0;\frac{d t_{TC,T}}{d x_{i}}={f_{r}s}_{r}y_{r}>0)$, while CRISPR-Cas-mediated gains are *decreasing* functions of TA activity *y* ($\frac{d t_{CT,C}}{d y_{i}}={{- x}_{r}s}_{i}\left( 1-f_{r} \right)<0;\frac{d t_{TC,C}}{d y_{r}}={{- x}_{i}s}_{r}\left( 1-f_{i} \right)<0)$: PSK destroys the advantages of competitor segregation, regardless of resident or invader role.

To investigate how increasing investment in CRISPR-Cas (*x*) impacts competitive outcomes (CRISPR gains versus TA gains), we analyse outcomes using differences in mutually exclusive transition paths Δ*_C_*, from a defined ecological context (*CT* or *TC*, Figure 1AB, equations E1-E4). To begin, we examine the ‘resident CRISPR-Cas’ context (*CT* cells), and define the resident CRISPR-Cas Δ*_Cr_* as

$\Delta_{Cr}= t_{CT,C}- t_{CT,T}= -s_{r}+ s_{i}x_{r}(1-y_{i}\left( 1+f_{i}-f_{r} \right))$	E5

When Δ*_Cr_* > 0, *CT* cells resolve to *C* cells more often, compared to *T* cells. When Δ*_Cr_* < 0, resolution to *T* cells dominates. We next examine how Δ*_Cr_* changes with increasing CRISPR-Cas investment, *x_r_*:

$\frac{d \Delta_{Cr}}{d x_{r}}=s_{i} (1-y_{i}\left( 1+f_{i}-f_{r} \right))$	E6

From equation E6 we can see that in the absence of invader TA activity (*y_i_* = 0), $\frac{d \Delta_{Cr}}{d x_{r}}=s_{i}$ reinforcing that resident CRISPR-Cas investments *x_r_* are always beneficial in absence of TA activity. When TA is active, we see that the return on investment ($\frac{d \Delta_{Cr}}{d x_{r}}$) is decreasing with *y_i_*. While decreasing, we note that given our assumptions *f_r_* ≥ *f_i_* and 0 ≤ *y_i_* ≤ 1, equation E6 cannot turn negative. For our default parameters (see parameterization table below), maximum *y_i_* = 0.66, *f_i_* = 0.04 and *f_r_* = 0.2, and therefore $y_{i}\left( 1+f_{i}+f_{r} \right)=0.66\times0.84=0.55$.

We next examine the context when the CRISPR-Cas plasmid is an invader (*TC* cells), and define the competitive outcome in this context Δ*_Ci_* as

$\Delta_{Ci}= t_{TC,C}- t_{TC,T}= -s_{i}+ s_{r}x_{i}(1-y_{r}\left( 1-f_{i}+f_{r} \right))$	E7

When Δ*_Ci_* > 0, *TC* cells resolve to *C* cells more often, compared to *T* cells. When Δ*_Ci_* < 0, resolution to *T* cells dominates. We next examine how Δ*_Ci_* changes with increasing CRISPR-Cas investment, *x_i_*:

$\frac{d \Delta_{Ci}}{d x_{i}}=s_{r} (1-y_{r}\left( 1-f_{i}+f_{r} \right))$	E8

From equation E8 we can again see that in the absence of TA activity (*y_r_* = 0), CRISPR-Cas is always beneficial $\frac{d \Delta_{Ci}}{d x_{i}}=s_{r}$. When TA is active, we see that the return on investment ($\frac{d \Delta_{Ci}}{d x_{i}}$) is decreasing with *y_r_*, and can turn negative if $y_{r}\left( 1-f_{i}+f_{r} \right)>1$. This condition is possible given our assumption that *f_r_* > *f_i_*, and is favoured by our assumption *y_r_* > *y_i_*. Indeed, we find this condition to be met for our default parameters. For these defaults, *y_r_* = 0.99, *f_r_* = 0.2 and *f_r_* = 0.2, and therefore $y_{r}\left( 1-f_{i}+f_{r} \right)=0.99\times1.16=1.15$.

Parametrization for graphical analyses.

In order to visualize our mathematical results in Figure 1, we set parameters as detailed in Table S1.

Table S1: Default parameter values and rationales for graphical analyses.

| Baseline segregation rate *s*; *s_i_* > *s_r_* | |
| --- | --- |
| *s_r_* = 1  *s_i_* = 1.49 | Segregation rate when resident is set at *s_r_* = 1 as a baseline. In comparison, *s_i_* adopts a higher value, as newly invaded single-copy plasmids are more likely to be lost by segregation than resident multi-copy plasmids. Large plasmids, which will be more likely to carry systems such as CRISPR-Cas and TA, are most frequently found at an average plasmid copy number (PCN) of 1.49 [3]. Note that setting *s_i_* = 1.49 is a conservative estimate, as gene expression (e.g. of stable segregation systems) together with copy number of newly invaded plasmids is lower compared to established plasmids [2]. |
| PSK benefit to nearby kin cells *f*; 1 > *f_r_* + *f_i_* > *f_r_* > *f_i_* ≥ 0 | |
| *f_r_* = 0.2  *f_i_* = 0.04 | As a baseline, *f_r_* captures the benefit resources freed up by post-segregational killing bring to neighbouring cells. *f_r_* = 0.2 assumes that PSK resources contribute to 20% as much growth of single-infected cells as loss of the competitor plasmid by segregation. The difference between *f_r_* and *f_i_* captures how well-structured an environment is, with *f_i_* = 1/5 * *f_r_*, we are assuming a dead cell is five times more likely to be surrounded by kin- than non-kin cells. A spatially structured environment is crucial to TA competitiveness [1]. |
| Strength of CRISPR-Cas *x*; *x_r_* > *x_i_* ≥ 1 | |
| *x_r_* = 1300  *x_i_* = 900  (max. values) | *x* describes the multiplicative effect CRISPR-Cas has on its competitor plasmid’s segregation rate (by cleaving it). In an analysis of plasmids targeted by a chromosomal CRISPR-Cas system, PCN of different plasmids was reduced to ~0.077-0.06% by CRISPR-Cas action depending on plasmid [4]. Thus, we estimate that CRISPR-Cas activity accelerates competitor segregation 1300-fold (≈ 1 / 0.00077). CRISPR-Cas activity on a newly invaded plasmid *x_i_* is scaled by PCN, leading to *x_i_* = 1/1.49 * *x_r_* ≈ 900. |
| Strength of TA *y*; 1 > *y_r_* > *y_i_* ≥ 0 | |
| *y_r_* = 0.99  *y_i_* = 0.66  (max. values) | We set the maximum *y_r_* value based on an average survival rate of post-segregational killing of 7 different TA systems of ~0.0067 [5]. We scaled *y_i_* by PCN, leading to *y_i_* = 1/1.49 * *y_r_* ≈ 0.66. |

**Supplementary Results 2 – Experimental results**

We assessed maintenance of vectors pCDF1b and pCDF1b_par by selectively plating mating mixes of *E. coli* hosts onto selective medium containing Streptomycin and comparing these with colony counts on non-selective medium (Figure S1). The proportion of Streptomycin-resistant colonies was never significantly different from 1 (T test followed by Bonferroni adjustment for multiple testing; Table 6). Additionally, when stamp plating, only two individual clones out of 2574 were found to be streptomycin sensitive, one of which was from a single-host control treatment. Vector presence was confirmed by PCR in all tested DH5α::SmR colonies.

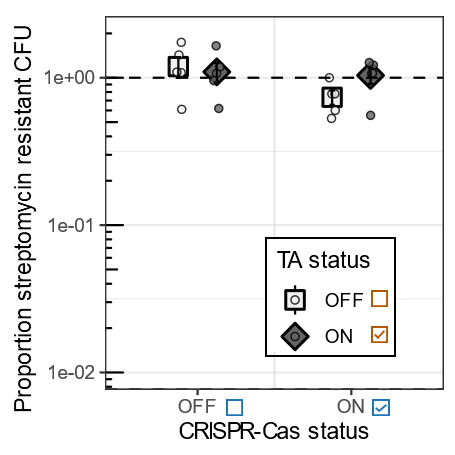

Figure S1: Mean ± standard error of the proportion of Streptomycin resistant Colony Forming Units (CFU) after mating was never significantly different from 1 (T test followed by Bonferroni adjustment for multiple testing; Table 6).

Three-strain competition shows putative generalisability of results

We repeated the two-strain competition experiment (Figure 2) using an additional plasmid host. In this way, we included an intermediate scenario where both plasmids invade an initially naïve, plasmid-free host, so that the mode of action of CRISPR-Cas could be either offensive or defensive. Here, we used DH5α::SmR as the initial CRISPR plasmid host, DH5α::CmR as the initial TA plasmid host, and DH5α::GmR as the naïve host (Figure S2B). A low limit of detection of TA plasmid RP4 in the defensive CRISPR-Cas host (Figure S2C) led to missing datapoints, especially where TA was turned off, and did not allow us to draw quantitative conclusions of the impact of TA status on the benefit conveyed by CRISPR-Cas in this host. Nonetheless, this analysis revealed that when CRISPR-Cas was acting defensively, the CRISPR-Cas plasmid pKJK5 won the competition in every case where CRISPR-Cas was switched on (Figure 2C; competitive ratio = 1.41 (N=1) and 1.94 ± 0.0085 respectively; significantly higher than 0 with p < 0.001 after T-test and Bonferroni adjustment for multiple testing with α = 0.005). In stark contrast, when CRISPR-Cas was acting offensively, switching on CRISPR-Cas had a differential effect depending on TA status of the competitor plasmid. When TA was absent from the competitor, switching on CRISPR-Cas brought little or no benefit to pKJK5 (Figure S2D; competitive ratio = 0.32 ± 0.13; p = 0.07). However, when the competitor plasmid’s TA was switched on, switching on CRISPR-Cas was clearly detrimental to pKJK5 (competitive ratio = -2.19 ± 0.11; p < 0.001).

In the naïve host, switching on CRISPR-Cas on its own benefitted pKJK5 (Figure S2E; competitive ratio 0.73 ± 0.1; p = 0.0019). When TA was switched on alongside this, there was no clear winner of the competition (competitive ratio = -0.21 ± 0.096; p = 0.1), however CRISPR-Cas still provided a benefit to pKJK5 relative to the scenario where CRISPR-Cas was switched off, where TA plasmid RP4 won the competition (competitive ratio = -0.94 ± 0.066; p < 0.001).

Overall, these data show that addition of a further plasmid host does not change the outcome of plasmid competition in hosts in which plasmids were already established.

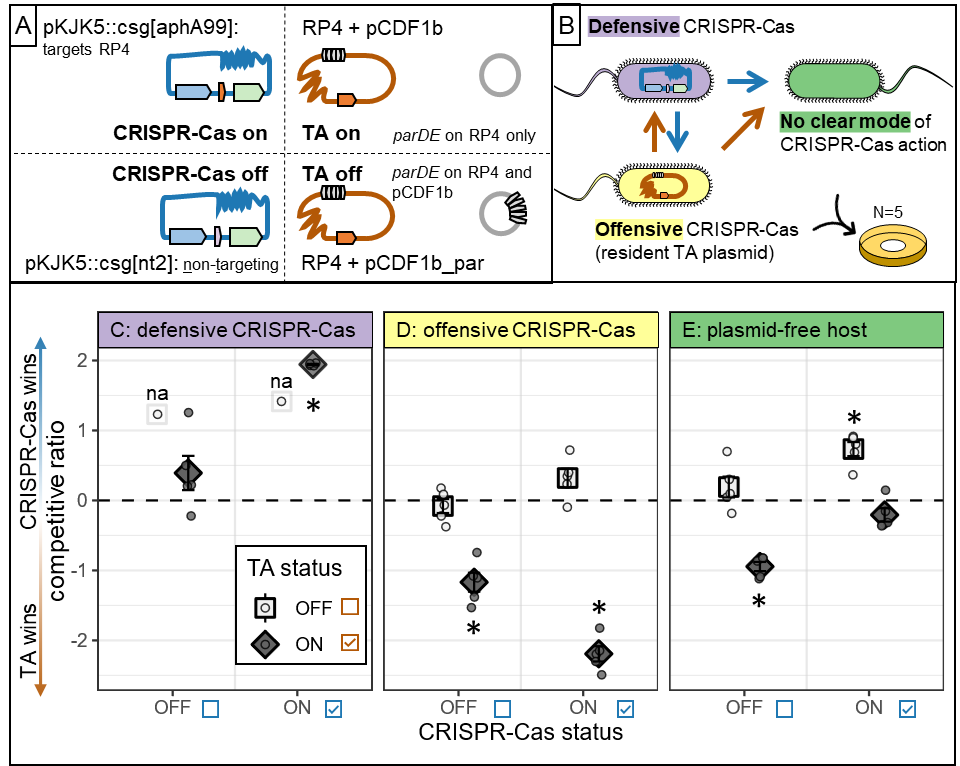

Figure S2: Plasmid competition results were upheld in a three-strain model system.

A: CRISPR-Cas and TA activity were tweaked on competitor plasmids in a binary manner by switching each system on or off. pKJK5::csg carries a gene cassette encompassing *cas9, sgRNA, GFP* with either an RP4-targeting or non-targeting guide, RP4 naturally carries TA system *parABCDE*. B: Plasmids were competed using three *E. coli* DH5α hosts with different plasmid content, allowing us to assess competitive outcome under defensive, offensive, or no clear mode of CRISPR-Cas action. After filter mating, plasmid content of each host was assessed by selective plating.

C-E: Mean ± standard error of the competitive ratio (log odds ratio of pKJK5-carrying hosts / RP4-carrying hosts) describes outcome of plasmid competition (N=5). Values >0 indicate CRISPR-Cas plasmid pKJK5 winning the competition, and values <0 indicate TA plasmid RP4 winning the competition for a certain host. The dashed line indicates a neutral outcome with competitive ratio = 0. Data are presented for treatments in which CRISPR-Cas and TA activity were toggled on or off in all combinations. Opacity of datapoints indicate number of replicates used to calculate means, this affects panel C TA-off data (N=1) and panel C TA-on & CRISPR-Cas-on data (N=4).

Stars indicate significant differences from 0 as assessed by T test and Bonferroni adjustment for multiple testing; *p < 0.005 with α = 0.005; see Table 3 for all p values. na – not assessed due to N=1 after removal of missing values (no carriage of TA plasmid RP4 recorded in 4 out of 5 replicates).

Competitive outcome is not due to RP4’s backbone

TA plasmid RP4 is a 60kB conjugative plasmid carrying many gene cassettes, including additional TA systems. Therefore, we sought to independently verify any causal fitness effects of *parABCDE* in plasmid competition. We constructed three vectors with different backbones of varying incompatibility group (IncP, pBR322, and IncQ). We engineered each vector to carry RP4’s *parABCDE* operon and competed these with CRISPR-Cas plasmid pKJK5 delivered to recipients using an *E. coli* DH5α::SmR donor. As non-TA carrying controls, we used the respective empty vector backbones. These vectors were not self-transmissible, therefore we measured the outcome of plasmid competition when CRISPR-Cas was acting offensively only, described here as the competitive ratio within DH5α::CmR recipients. Firstly, the outcome of plasmid competition showed that previously observed results of competition with RP4 were largely upheld in this model system using a different DH5α tag variant as the CRISPR-Cas plasmid host, and also when lacking a bystander plasmid: switching CRISPR-Cas on in the presence of TA was detrimental to pKJK5 in offense (Figure S3A-B; p < 0.05 after fitting a GLM and a linear model, see Methods). When competing pKJK5 with vectors of varying incompatibility groups, we found that the TA system hindered the offensive ability of CRISPR-Cas on pKJK5. The competition outcome with pOGG99 (an IncP vector) directly mirrored our results recorded with RP4: turning on CRISPR-Cas in the absence of TA benefitted pKJK5, while turning on CRISPR-Cas in the presence of TA was detrimental to the CRISPR-Cas plasmid (Figure S3C; p < 0.001 and p = 0.47 in the absence and presence of TA respectively after fitting a GLM and carrying out Tukey’s post hoc test, see Methods). For the remaining two vectors (pBR322 backbone vector pHERD99 and IncQ backbone vector pSEVA251-99), the competitive ratios followed the same trend, and we observed a significant increase following CRISPR-Cas expression in favour of pKJK5 in the absence of TA for pHERD99 (Figure S3D, p < 0.001 after fitting a GLM and Tukey’s post hoc test), and a significant decrease, showing a detrimental CRISPR-Cas system, in the presence of TA for pSEVA251-99 (Figure S3E, p < 0.001 after fitting a GLM and Tukey’s post hoc test). The remaining comparisons between competitive ratios remained non-significant, but crucially CRISPR-Cas was never detrimental in the absence of TA, and never beneficial in the presence of TA.

The data generated with these simplified model systems strongly support our conclusion that RP4’s *parABCDE* operon limits CRISPR-Cas effectivity.

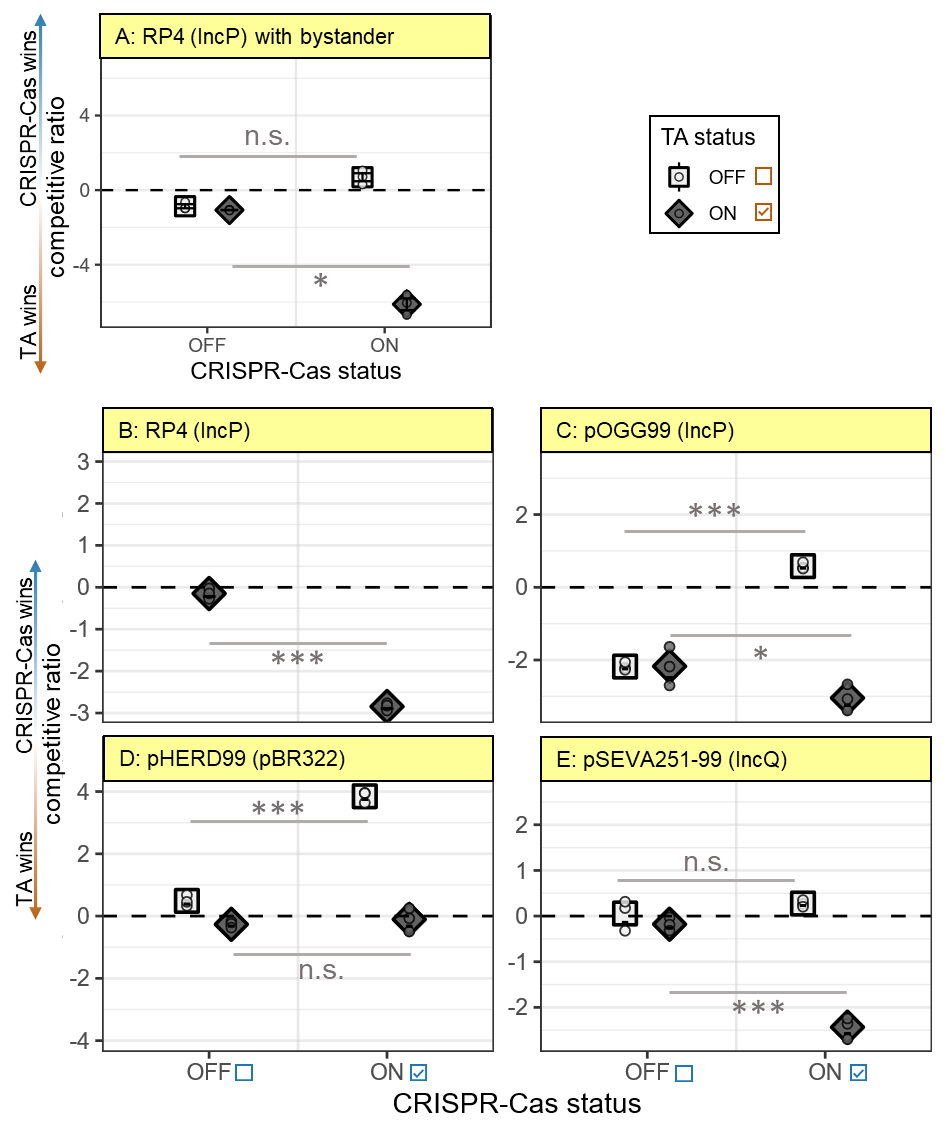

Figure S3: Outcome of competition with cloning vectors. Mean ± standard error of the competitive ratio (log10 of pKJK5-carrying hosts / competitor-carrying hosts) describes the outcome of plasmid competition (N=5). Values <0 indicate CRISPR-Cas plasmid pKJK5 winning the competition, and values >0 indicate the TA competitor plasmid winning the competition for a DH5α host where CRISPR-Cas was acting offensively. Data are presented for treatments in which CRISPR-Cas and TA activity were toggled on or off in all combinations. RP4 competition outcome (A-B) in this model system matched the outcome observed earlier (Figure 2B). Relative differences between treatments are indicated with grey lines and were assessed by Tukey’s HSD after fitting individual Generalised Linear Models; see methods and Table 5-6 for model details and p values. *p<0.05; ***p<0.001; n.s. not significant p>0.63.

Throughout experimental work, RP4 carriage was assessed by selection using kanamycin, resistance to which is conferred by the CRISPR-Cas9 target gene *aphA*. This means theoretically, RP4 escape mutants with non-functional *aphA* genes could skew RP4 carriage results. However, plasmid escape of CRISPR-Cas by target mutation is unusual [6]. We carried out a control experiment in which we mated strains exactly as in Figure S2AB, with the exception of omission of bystander plasmid pCDF1b (i.e. TA was active in all treatments). We confirmed that the outcome of RP4 carriage was largely robust to altering the selective agent to ampicillin (resistance is conferred by *blaTEM* on the opposite side of *aphA* on RP4, >25kB away). RP4 carriage was assayed as moderately lower when plating on kanamycin (10.11±2.68 – 13.44±3.69%) than when plating on ampicillin (42.72±6.0% – 68.64±16.75% for non-targeting and targeting treatments respectively) in only the DH5α::GmR host (Figure S4C), but this was observed for both pKJK5::csg treatments (Figure S4). Therefore, these discrepancies are likely due to discrepancies in resistant colonies when using different combinations of antibiotics, and we infer that RP4 escaping pKJK5::csg[aphA99] targeting by *aphA* mutation is negligible in this work.

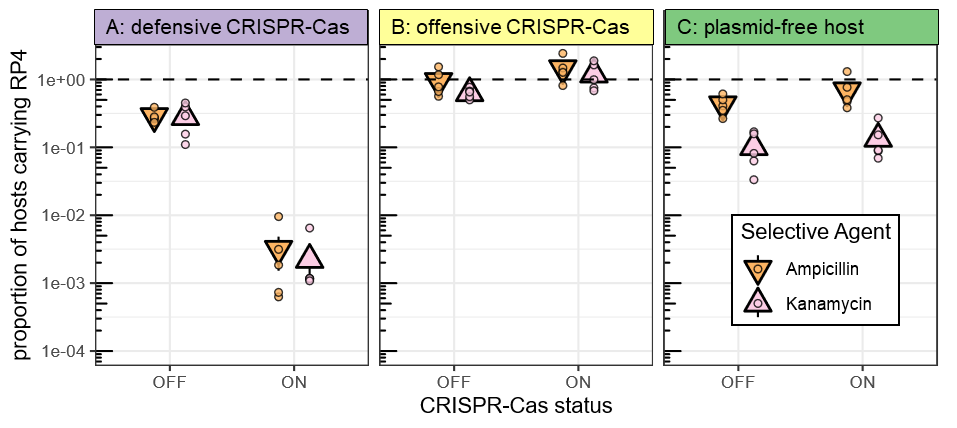

Figure S4: Assessed RP4 proportion is largely robust to antibiotic used to select for RP4.

Mean and standard error of RP4 content of hosts in a control mating experiment lacking a bystander plasmid assessed by ampicillin or by kanamycin, N=5. Data are presented for CRISPR-Cas switched on or off.

**Supplementary Results 3 – Bioinformatics Results.**

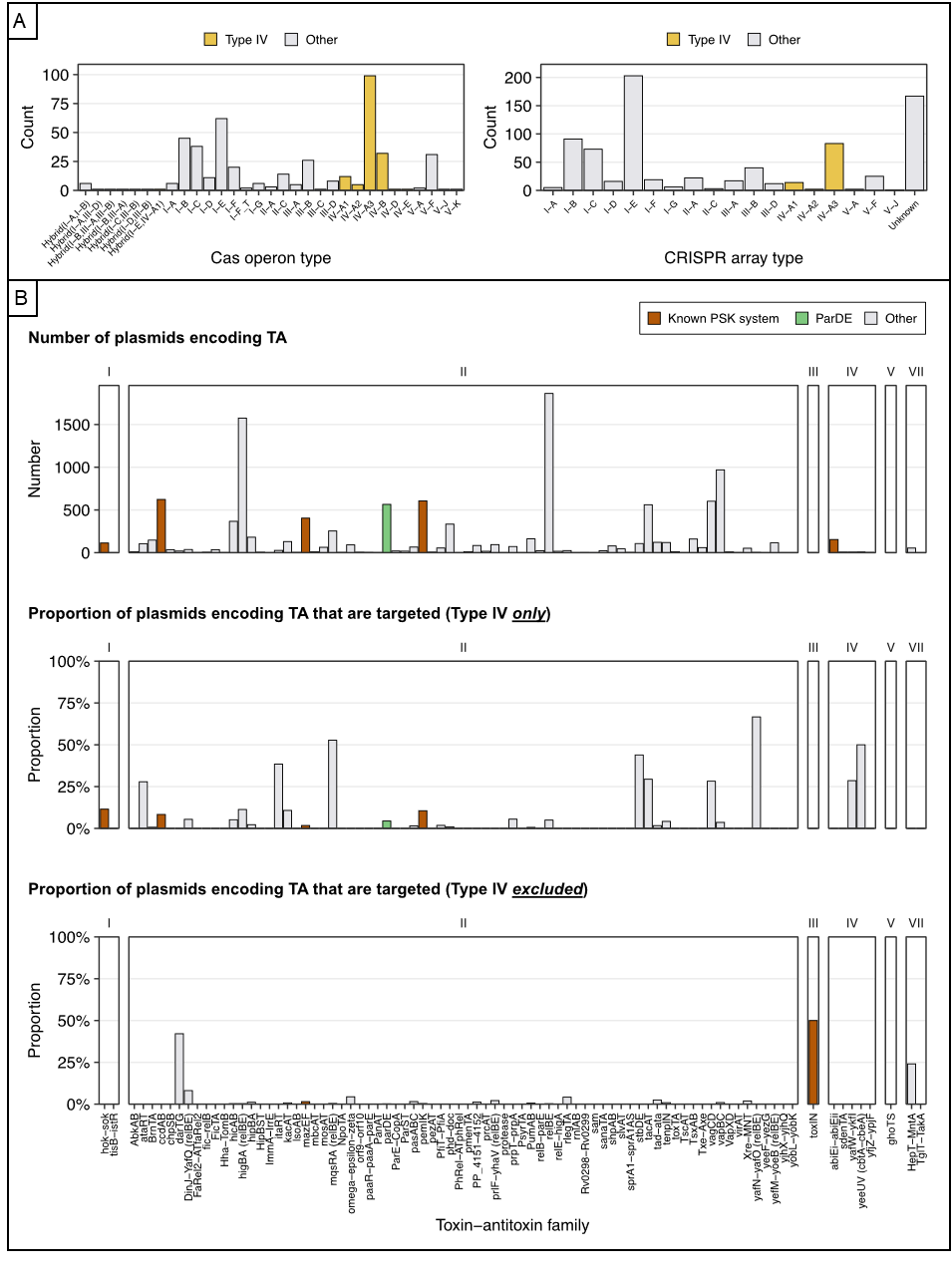

Figure S5: Plasmid-borne CRISPR-Cas systems and their targets.

A: CRISPR-Cas systems encoded on plasmids; split into distribution of Cas operon and CRISPR array types. Type IV systems are highlighted to emphasize their predominance

B: The total number of plasmids encoding each TA family, highlighting *parDE* and other well-studied post-segregational killing (PSK) systems [7]. The other bar charts show the proportion of plasmids encoding each TA family targeted by plasmid borne Type IV and non-Type IV CRISPR-Cas systems.

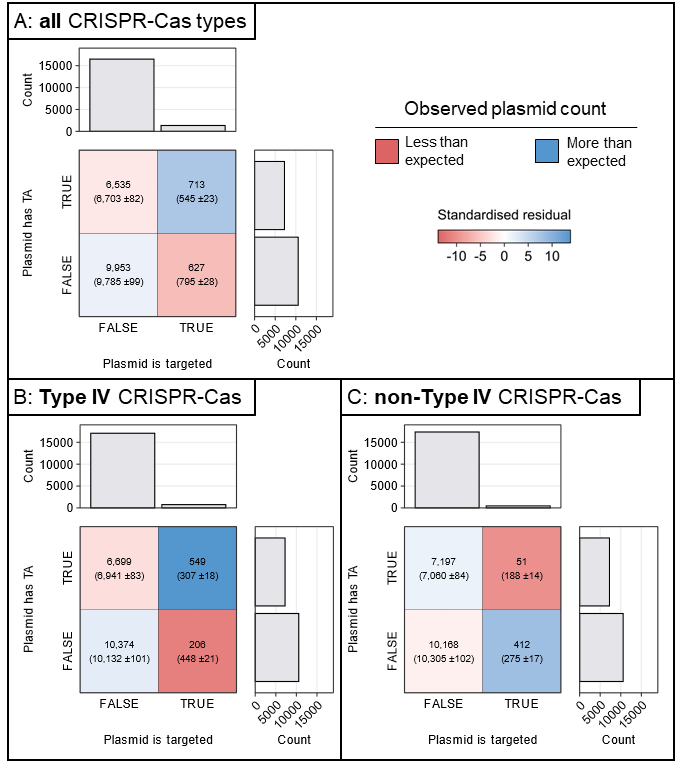

Figure S6: Contingency tables of expected versus observed values for the association between plasmid-borne CRISPR-Cas targeting and TA carriage.

Contingency tables showing the relationship between plasmid targeting and TA presence. The tables display the observed counts of plasmids with or without TA systems and whether they are targeted or not by CRISPR spacers in another plasmid, either for all CRISPR-Cas types (A), Type IV CRISPR-Cas only (B), or for non-Type IV CRISPR-Cas (C). The bar plots show the sum of counts for each vertical or horizontal group. Facets in the table are coloured by standardised residual after Pearson’s χ^2^ test, indicating magnitude and directionality of association (red – less than expected, blue – more than expected).
